# Differential memory enrichment of cytotoxic CD4 T cells in Parkinson’s disease patients reactive to α-synuclein

**DOI:** 10.1101/2025.04.21.649867

**Authors:** Antoine Freuchet, Emil Johansson, April Frazier, Irene Litvan, Jennifer G Goldman, Roy N Alcalay, David Sulzer, Cecilia S. Lindestam Arlehamn, Alessandro Sette

## Abstract

Parkinson’s disease (PD) is a complex neurodegenerative disease with a largely unknown etiology. Although the loss of dopaminergic neurons in the substantia nigra pars compacta is the pathological hallmark of PD, neuroinflammation also plays a fundamental role in PD pathology. We have previously reported that PD patients have increased frequencies of T cell reactive to peptides from *α*-synuclein (*α*-syn). However, not all PD participants respond to *α*-syn. Furthermore, we have previously found that CD4 T cells from PD participants responding to *α*-syn (PD_R) are transcriptionally distinct from PD participants not responding to *α*-syn (PD_NR). To gain further insight into the pathology of PD_R participants, we investigated surface protein expression of 11 proteins whose genes had previously been found to be differentially expressed when comparing PD_R and healthy control participants not responding to *α*-syn (HC_NR). We found that Cadherin EGF LAG seven-pass G-type receptor 2 (CELSR2) was expressed on a significantly higher proportion of CD4 effector memory T cells (T_EM_) in PD_R compared to HC_NR. Single-cell RNA sequencing analysis of cells expressing or not expressing CELSR2 revealed that PD_R participants have elevated frequencies of activated T_EM_ subsets and an almost complete loss of cytotoxic T_EM_ cells. Flow cytometry analyses confirmed that Granulysin^+^ CD4 cytotoxic T_EM_ cells are reduced in PD_R. Taken together, these results provide further insight into the perturbation of T cell subsets in PD_R, and highlights the need for further investigation into the role of Granulysin^+^ CD4 cytotoxic T_EM_ in PD pathology.

## Introduction

Parkinson’s disease (PD) is a progressive neurodegenerative disease affecting 10 million people worldwide, which is increasing with age^1^. It is characterized by the loss of dopaminergic (DA) neurons in the substantia nigra (SN)^2^, and the accumulation of abnormal aggregation of misfolded *α*-synuclein (*α*-syn), called Lewy bodies, in neurons in the brain stem as well as other brain and nervous system regions^3^. While the number of DA neurons decreases and Lewy bodies accumulate, motor symptoms such as tremor, rigidity and postural instability appear and progress^4^. Additionally, non-motor symptoms (e.g., sleep disorders, gastrointestinal dysfunction) can manifest up to 20 years before PD diagnosis and other non-motor symptoms (e.g., cognitive impairment) occur during the course of disease^5^. Today’s challenges of early diagnosis and symptomatic treatments are limited to a few options because of the complexity and heterogeneity observed and described in PD^6^. Thus, further work is needed to unravel and better comprehend the disease with an ultimate goal to develop targeted therapies.

Neuroinflammation is a common feature in PD^7^. Reports have shown activated microglia in the SN of PD brain, a characteristic of inflammation^8^. By secreting pro-inflammatory cytokines, activated microglia can directly induce neurotoxicity, disrupt the blood-brain barrier (BBB)^9^, or recruit immune cells from the periphery to the central nervous system (CNS). Indeed, neuropathological analysis of PD brains demonstrated increased T cell infiltrations in the substantia nigra^10–12^. More importantly, PD has an autoimmune component represented by an increased reactivity to *α*-syn by T cells from the periphery^13, 14^, indicating promise for immune-based therapies. However, not all PD patients are reactive to *α*-syn (PD_NR; non-responder)^15^, confirming the heterogeneity of the disease and suggesting the existence of other neuroantigens^16^. Interestingly, transcriptomes of CD4 and CD8 memory T cells from PD_R (*α*-syn responders) are different from those from non-responders. Pathways analysis on upregulated genes by CD4 and CD8 memory T cells from PD_R revealed specific enrichment for PD, oxidative phosphorylation, and inflammation pathways^15^, highlighting *α*-syn responders as a subgroup of PD. Thus, we hypothesized that we can identify membrane target(s) specific to PD_R using their gene signature, and further investigate the heterogeneity at the single-cell level of PD participants reactive to *α*-syn.

Here, using a bulk RNAseq dataset to identify candidates, we show in PD_R that Cadherin EGF LAG seven-pass G-type receptors 2 (CELSR2) protein is expressed by T cells and specifically enriched in CD4 T cell effector memory (T_EM_). We thereafter compared the single-cell transcriptome between CELSR2^+^ and CELSR2^-^ T_EM_ cells in PD_R and HC_NR participants, and found an enrichment of activated and differentiating cells in CELSR2+ PD_R samples. Strikingly, we also discovered an almost complete lack of CD4 cytotoxic T cells cluster in CD4 T_EM_ in PD_R compared HC_NR (Healthy controls non-responders to *α*-syn). Using flow cytometry, we confirmed that Granulysin^+^ CD4 cytotoxic T_EM_ cells are reduced in PD_R, a finding that promises to open new research avenues.

## Results

### CESLR2 expression is significantly higher on CD4 effector memory T cells in PD participants responding to *α*-syn compared to healthy controls

Previous studies showed that *α*-syn T cell reactivity is associated with PD^13^, and that this inflammatory adaptive immune response is influenced by disease evolution and other factors^14^. Accordingly, PD patients can be classified as a function of *α*-syn T cell reactivity in PD_R and PD_NR^15^, highlighting *α*-syn responders as a subgroup of PD. This classification allowed us to detect specific differences in the transcriptomes of CD4 memory T cells between PD_R and PD_NR^15^.

Here, we undertook a more detailed analysis of a previously published bulk RNAseq dataset associated with CD4 memory T cells (T mem) from healthy controls who do not respond to *α*-syn (HC_NR), PD_NR, and PD_R participants^15^. We extracted the list of genes expressed at significantly higher levels in PD_R compared to PD_NR or HC_NR. These genes were filtered for protein-coding genes, followed by proven or predicted membrane localization according to the Human Protein Atlas, and whether commercial specific antibodies were available.

We obtained 11 potential candidates (**Table 1**): *LSMEM1, KCNH4, RNF152, CD300LB, AIG1, APOL1, ABCD2, CELSR2, LMO7, ABCC3* and *FFAR3*. Five (*LSMEM1, AIG1, APOL1, ABCD2, and CELSR2)* were upregulated in PD, while the remaining genes were downregulated in PD. As the correlation between gene and protein expression is often imperfect^17^, we next performed flow cytometry assays to examine which differentially expressed genes also differed significantly at the protein level in memory CD4 T cells (total CD45RA^+^CCR7^-^, CD45RA^-^CCR7^+^, and CD45RA^-^ CCR7^-^) from HC_NR (n=12-17) and PD_R (n=12-17) (**Supplemental Figure 1A, 1B**). Donors were selected based on sample availability, and in line with the increased risk of males to develop PD^18^, we had a significantly higher proportion of male PD_R participants compared to HC_NR (p=0.001, **Supplemental Data 1**). We found that the frequency of CELSR2 expressing cells were significantly higher in PD_R compared to HC_NR memory CD4 T cells (HC_NR, median=46.9%; PD_R, median=70.8%; p=0.024), while no difference was found for naïve CD4 T cells (**Figure 1A, 1B**). We further investigated CELSR2 expression in memory vs naïve (CD45RA^+^CCR7^+^) CD4 T cells. Among both HC_NR and PD_R, CELSR2 positive cells were significantly enriched in memory compared to naïve CD4 T cells (**Figure 1A**), and similarly the CELSR2 mean fluorescent intensity (MFI) was found to be higher in CELSR2 expressing memory compared to naïve CD4 T cells (**Supplemental Figure 1C**). Within memory CD4 T cell subsets, a higher frequency of CELSR2^+^ was found in T_EM_ (CD45RA^+^CCR7^+^), compared to central memory (T_CM_; CD45RA^-^CCR7^+^), (p<0.0001) and effector memory T cells re-expressing CD45RA (T_EMRA_; CD45RA^+^CCR7^-^) (p<0.0001) (**Figure 1C**), but no difference in CELSR2 MFI was found (**Supplemental Figure 1D**). Additionally, we observed a significantly greater frequency of CELSR2^+^ CD4 T_EM_ in PD_R compared to HC_NR (p=0.0216; **Figure 1C**).

**Figure 1.**
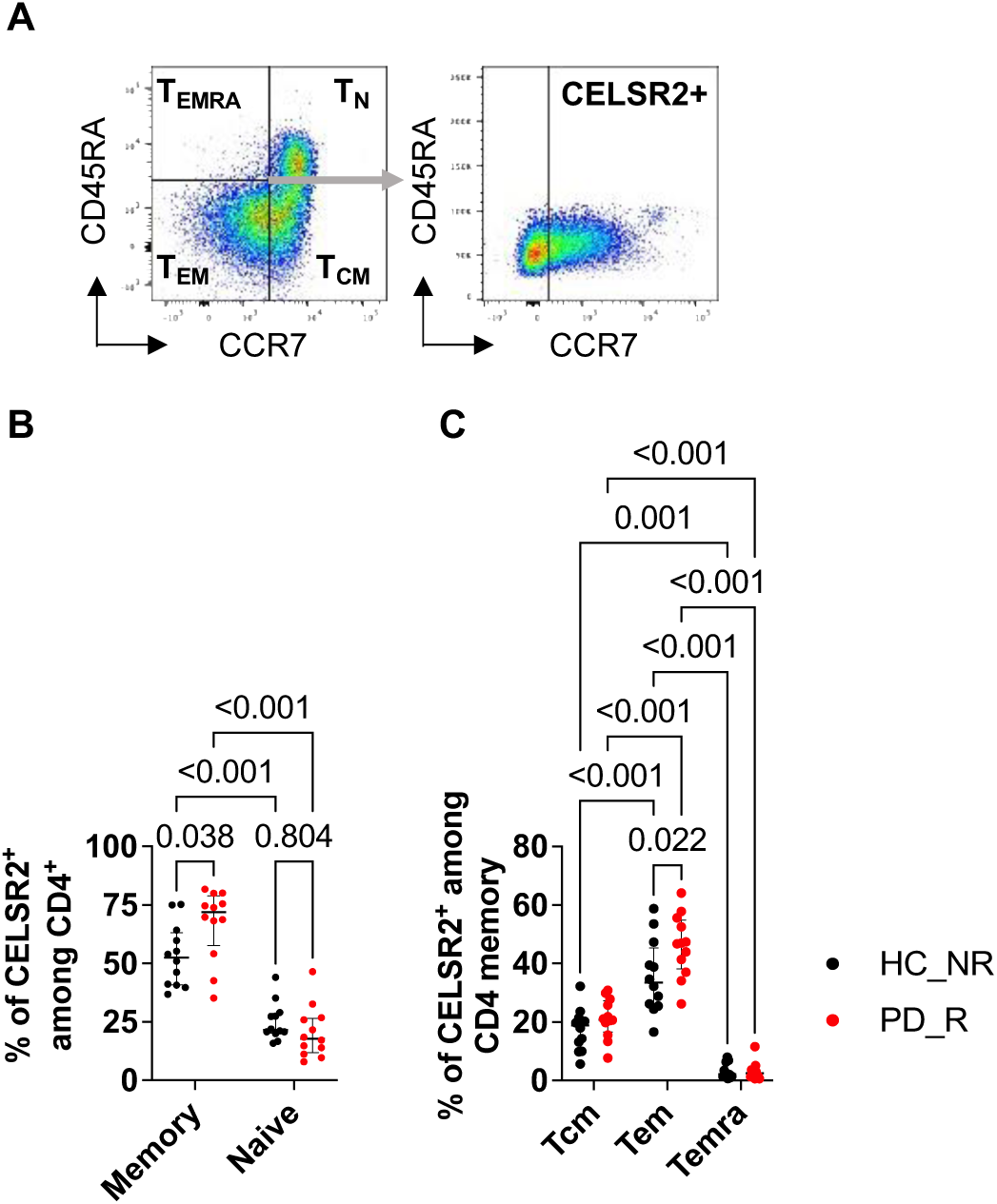
CELSR2^+^ cells are enriched in CD4 T_EM_ in PD_R participants. **(A)** Representative gating strategy of how CELSR2+ CD4 T cells were identified in four different memory subpopulations in HC_NR (n=12) and PD_R (n=12) participants. Frequency of CELSR2+ cells were identified for each of the four subpopulations. Complete gating strategy is displayed in **Supplemental** Figure 1. **(B)** Frequency of memory (combined central memory (T_CM_): CD45RA^-^ CCR7^+^; effector memory (T_EM_): CD45RA^-^CCR7^-^; and effector memory T cells re-expressing CD45RA (T_EMRA_): CD45RA^+^CCR7) vs. naïve T cells (T_N_: CD45RA^+^CCR7^+^) among CELSR2+ cells. Results are represented as median with interquartile range. Mann Whitney U test. **(C)** Frequency of T cell memory subpopulations (T_CM_, T_EM_, and T_EMRA_) among memory CELSR2^+^ CD4 T cells. Results are represented as median with interquartile range. 2way ANOVA and Tukey’s multiple comparisons test.

**Table 1.** Protein candidates screened for T cell expression.

| Gene | Gene type | Cell localization<br>(Protein Atlas) | PD_R vs PD_NR |  | PD_R vs HC_NR |  |
| --- | --- | --- | --- | --- | --- | --- |
|  |  |  | log2 FC | adj p-value | log2 FC | adj p-value |
| *LSMEM1* | Protein coding | Membrane | 1.63 | 6.9E-08 | 1.37 | 6.8E-6 |
| *KCNH4* | Protein coding | Membrane | -1.08 | 5.3E-46 | -1.95 | 5.2E-43 |
| *RNF152* | Protein coding | Intracellular,<br>Membrane | -0.98 | 4.8E-50 | -1.13 | 3E-38 |
| *CD300LB* | Protein coding | Intracellular,<br>Membrane | -0.92 | 2.3E-17 | -0.87 | 7.8E-14 |
| *AIG1* | Protein coding | Intracellular,<br>Membrane | 1.7 | 3.1E-5 | 1.25 | 6.9E-3 |
| *APOL1* | Protein coding | Membrane,<br>Secreted | 1.67 | 1.1E-4 | 1.95 | 1.3E-4 |
| *ABCD2* | Protein coding | Membrane | 1.87 | 1.7E-4 | 1.56 | 1.2E-3 |
| *CELSR2* | Protein coding | Membrane | 1.16 | 9.9E-4 | 1.67 | 8.5E-5 |
| *LMO7* | Protein coding | Intracellular,<br>Membrane | 1.41 | 1.4E-2 |  |  |
| *ABCC3* | Protein coding | Intracellular,<br>Membrane | -1.61 | 8.6E-40 | -0.72 | 1.9E-26 |
| *FFAR3* | Protein coding | Membrane | -0.88 | 1E-17 | -0.59 | 1.8E-18 |

Altogether, these experiments demonstrated that CESLR2 is expressed on the surface of various T cell subsets. The experiments further pinpointed CD4 T_EM_ as a particular T cell subset associated with increased frequency of CESLR2 expressing cells in PD_R compared to HC_NR.

### Differential frequency CD4 memory T cells as a function of PD status and CELSR2 positivity

Little is known about the heterogeneity associated with CELSR2 expression in T cells. Therefore, we next sorted CD4 T_EM_ CELSR2^+^ and CELSR2^-^ from PD_R (n=4; Female=2, Male=2) and HC_NR (n=4; Female=0, Male=4, p=0.429) (**Figure 2A, Supplemental Figure 1E, Supplemental Data 2**), and analyzed them using single-cell RNA sequencing. After filtering the data based on standard quality control pipelines, a shared nearest neighbor (SNN) based clustering algorithm revealed nine different clusters (**Supplemental Figure 2**). The cluster were subsequently named based on their top 50 expressed genes (**Figure 2B; Supplemental Figure 3, Supplemental Data 3**). *Cluster 0 – RP^+^* T_EM_ gene signature contains 32/50 ribosomal protein genes. Genes known to be important for regulation of immune responses, interaction with antigen presenting cells, and differentiation (NR4A3, *FHIT*, *SLC9A9,* SEMA4A) were highly expressed in *Cluster 1 – Differentiating* T_EM_. Clusters 2 and 3 have been defined as *T_H_1* T_EM_ and *T_H_17* T_EM_, respectively, as their gene signature includes *GZMK, CCL5, DUSP2, KLRB1, ITGA4* and *IL4I1, KLRB1, CCR6,* respectively. *Cluster 4 – Cytotoxic* T_EM_ was associated with high expression of several genes associated with cytotoxicity, such as *GZMB, GZMH, GNLY, NKG7, CCL4, ZNF683, KLRD1, PRF1, CST7, CCL5,* and *GZMA. Cluster 5* was denominated as *T_reg_* T_EM_ because it is associated with a high level of expression of *IKZF2, IL2RA, CTLA4,* and *TIGIT*. Several *HLA* genes (*HLA-DRA, HLA-DRB1, HLA-DPA1, HLA-DQB1, HLA-DPB1*) expressed in *Cluster 6* are associated with activation^19^, which was hence named *Activated* T_EM_. The top gene of *Cluster 7 – EDA^+^* T_EM_ is *EDA* (log2FC=4.6), a TNF family ligand with an unknown function in T cells^20^. *BCL6*, the key transcription factor for follicular T helper cells (T_FH_)^21^, was the top gene in *Cluster 8 – T_FH_* T_EM_.

**Figure 2.**
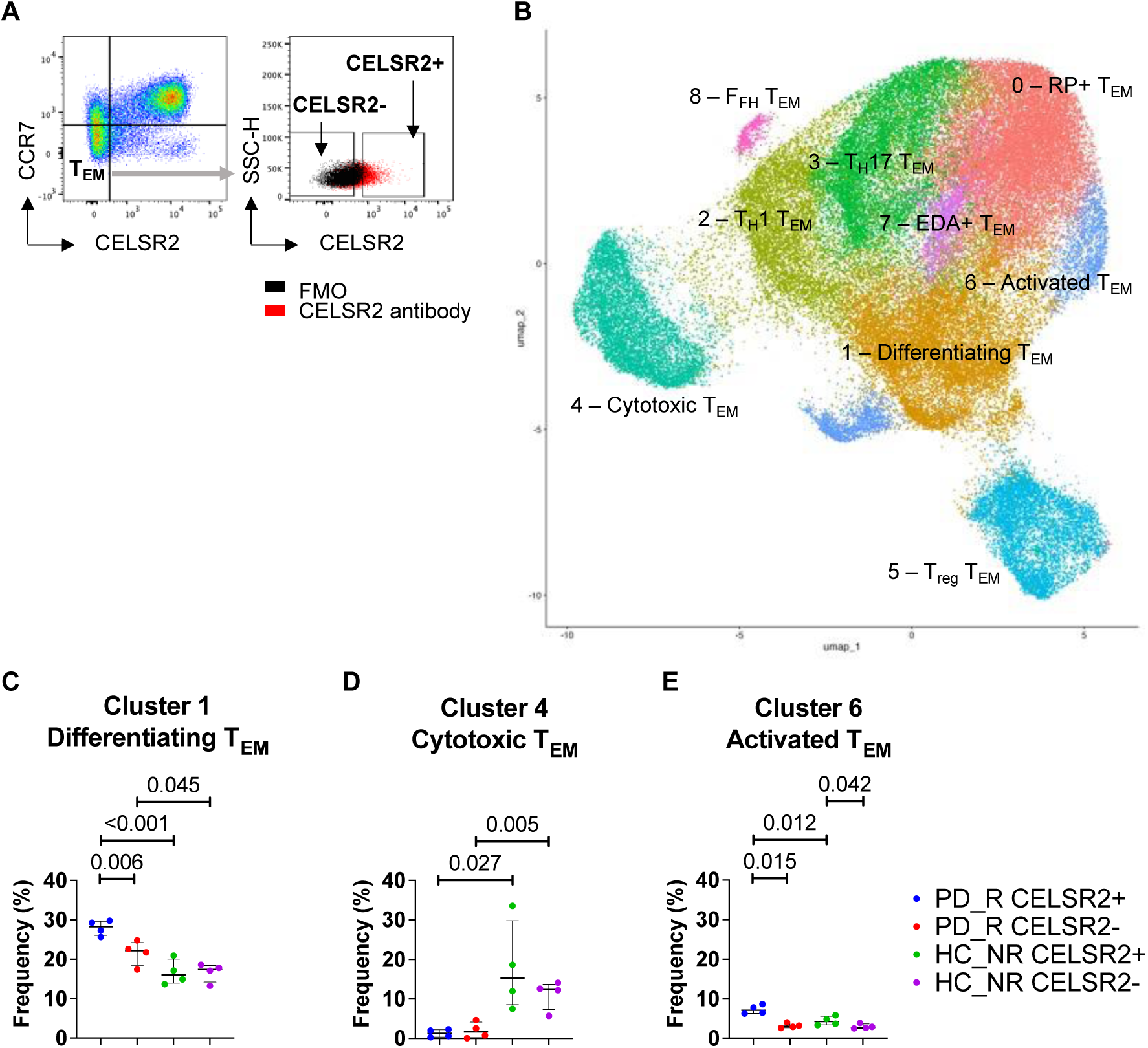
scRNAseq of CD4 T_EM_ highlights distribution differences specific to CELSR2 expression and/or PD_R. **(A)** Representative gating strategy for how CELSR2^+^ and CELSR2^-^ T_EM_ cells were identified and sorted from HC_NR (n=4) and PD_R (n=4) participants for subsequent scRNAseq analysis. **(B)** UMAP of CD4 T_EM_ CELSR2^-^ and CESLR2^+^ T cells. Cluster frequencies of Cluster 1 – Differentiating T_EM_ **(C)**, Cluster 4 – Cytotoxic T_EM_ **(D),** and Cluster 6 – Activated T_EM_ **(E)** were analyzed and compared between HC_NR and PD_R in CD4 T_EM_ expressing CELSR2 or not. Results are represented as median with interquartile range. Frequencies were compared using paired t test within PD_R or HC_NR samples, and unpaired t test to compare frequency between PD_R and HC_NR subgroups.

Next, we analyzed the distributions of each cluster across the different participant cohorts. Significant differences were seen in the case of Cluster 1 – Differentiating T_EM_, which was enriched in PD_R versus HC_NC samples, regardless of the CELSR2 positive or negative status (**Figure 2C**). The largest difference in cluster frequency was observed in PD_R CELSR2^+^ compared to HC_NR CELSR2^+^ (PD_R, median=28.3%; HC_NR, median=16.0%; p<0.001). Less prominent differences were also found comparing PD_R CELSR2^-^ vs HC_NR CELSR2^-^ (PD_R, median=22.2%; HC_NR, median=17.5%; p=0.045), as well as CESLR2^+^ vs CESLR2^-^ PD_R participants (CESLR2^+^, median=28.3%; CESLR2^-^, median=22.2%; p=0.006). Interestingly, a striking decrease in the frequency of Cluster 4 – Cytotoxic T_EM_ in PD participants was observed, both in the CELSR2^+^ (PD_R, median=1.3%; HC_NR, median=15.3%; p=0.027) and the CELSR2^-^ (PD_R, median=1.6%; HC_NR, median=12.4%; p=0.005) samples (**Figure 2D**). The frequency of Cluster 6 - Activated T_EM_ was the only cluster found to be enriched in CELSR2^+^ samples in both PD and HC participants (**Figure 2E).** Specifically, this cluster is enriched in CELSR2^+^ compared to CELSR2^-^ PD_R samples (CELSR2^+^, median=7.1%; CELSR2^-^, median=3.1%; p=0.015), in CELSR2^+^ compared to CELSR2^-^ HC_NR samples (CELSR2^+^, median=4.3%; CELSR2^-^, median=2.8%; p=0.042), as well as in PD_R CELSR2^+^ compared to HC_NR CELSR2^+^ (PD_R, median=7.1%; HC_NR, median=3.4%; p=0.012). However, no difference was found between CELSR2^-^ PD_R and HC_NR samples. We noted no significant difference between PD_R and HC_NR in the frequency of the other subsets. ***(Supplemental Fig 4*).**

Thus, single-cell analysis of CELSR2^+^ and CELSR2^-^ CD4 T_EM_ in PD_R and HC_NR reveals distribution differences in three CD4 T_EM_ clusters.

### CELSR2 expression is associated with few transcriptomic changes

We next determined the impact of CELSR2 expression on the RNA transcriptome in Cluster 1 – Differentiating T_EM_, Cluster 4 – Cytotoxic T_EM_, and Cluster 6 – Activated T_EM_, as the frequency of these clusters had been found to differ between CELSR2^+^ and CELSR2^-^ samples. In PD_R participants, 41 genes were found to be differentially expressed in Cluster 1 – Differentiating T_EM_ (**Figure 3A**), including elevated expression of *HLA-DRA*, *HLA-DQA2*, *HLA-DQA1*, *HLA-DRB1*, and *IL9R* in CELSR2^+^ cells, which were significantly enriched in the biological process Immunoglobulin Mediated Immune Response (**Supplemental Data 4**), suggesting a potential increased activation of these cells. In contrast, zero and one genes were differentially expressed in between CELSR2^+^ and CELSR2^-^ cells from Cluster 4 – Cytotoxic T_EM_ and Cluster 6 – Activated T_EM_, respectively (**Supplemental Figure 5A, 5B**). In HC_NR participants, 57 genes were differentially expressed between CELSR2^+^ and CELSR^-^ cells in Cluster 4 – Cytotoxic T_EM_ (**Figure 3B**), while six and one genes were differentially expressed in Cluster 1 – Differentiating T_EM_ and Cluster 6 – Activated T_EM_, respectively (**Supplemental Figure 5C, 5D**). Although no biological processes were enriched among the differentially expressed genes in HC_NR cells, we noted that genes associated with cytotoxic CD4 T cells, such as *GZMB*, *KLRD1*, *FCGR3A*, and *GNLY*, were among the upregulated genes in CELSR2^+^ cells. We therefore took a targeted approach and created a module score based on the expression of 109 genes that were both detected in this dataset, and included in the biological process Leukocyte mediated cytotoxicity (**Supplemental Data 5**). We found that the Leukocyte mediated cytotoxicity gene signature was enhanced in CELSR2^+^ compared to CELSR^-^ HC_NR participant cells (**Figure 3C**).

**Figure 3.**
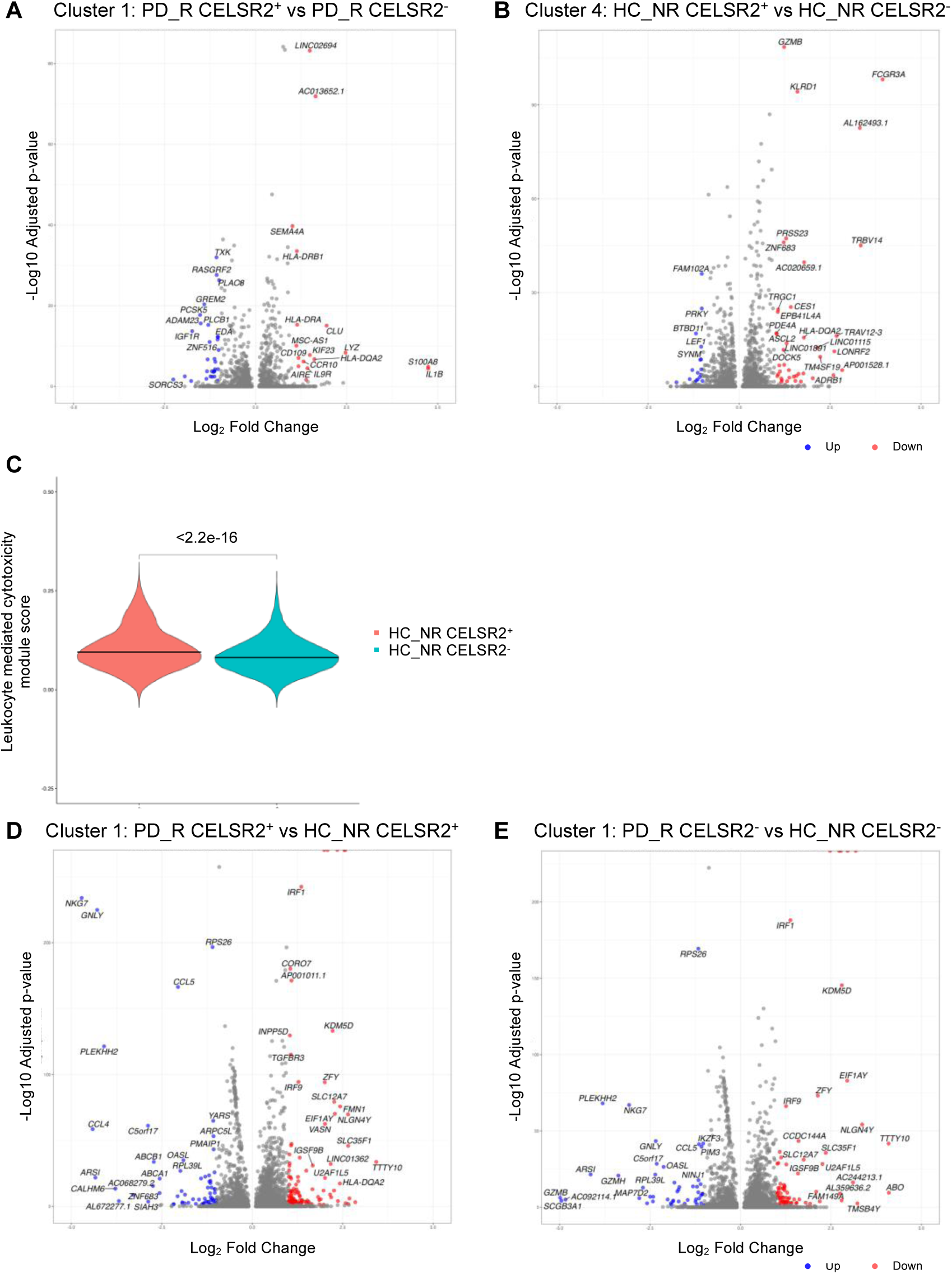
Overall expression of CELSR2 does not significantly impact transcriptomics, yet correlates with an enhanced cytotoxic gene signature in Cluster 4 – Cytotoxic T_EM_ in HC_NR. Volcano plots comparing differential gene expression (DEG) of PD_R CELSR2^+^ vs PD_R CELSR2^-^ in Cluster 1 – Differentiating T_EM_ **(A)** and HC_NR CELSR2^+^ vs HC_NR CELSR2^-^ in Cluster 4 – Cytotoxic T_EM_ **(B)**. Red points are significantly upregulated in CELSR2^+^samples and blue points are significantly higher in CELSR2^-^ samples (adjusted p value < 0.05 and Log_2_ fold change >1). (**C**) Visualization of Leukocyte mediated cytotoxicity module score in CELSR2^+^ and CELSR2^-^ cells from HC_NR participants. Data are represented as median, and p value obtained using an unpaired t test. The score was created using genes both detected in this study and included in the Leukocyte mediated cytotoxicity biological process (GO:0001909). Volcano plots comparing differential gene expression (DEG) of PD_R CELSR2^+^ vs HC_NR CELSR2^+^ cells **(D)** and PD_R CELSR2^-^ vs HC_NR CELSR2^-^ cells in Cluster 1 – Differentiating T_EM_ **(E)**. Red points are significantly upregulated in PD_R^+^ cells and blue points significantly higher in HC_NR (adjusted p value < 0.05 and Log_2_ fold change >1).

We next examined if cells from Cluster 1 – Differentiating T_EM_, Cluster 4 – Cytotoxic T_EM_, and Cluster 6 – Activated T_EM_ in PD_R and HC_NR participants displayed transcriptional differences. The number of differentially expressed genes were higher between PD_R and HC_NR for CELSR2^+^ compared to CELSR2^-^ cells for from Cluster 1 – Differentiating T_EM_ (198 and 129, respectively), Cluster 4 – Cytotoxic T_EM_ (164 and 129, respectively), and Cluster 6 – Activated T_EM_ (86 and 42, respectively) (**Figure 3D, 3E**). Among the differentially expressed genes in Cluster 1 – Differentiating T_EM_, PD_R CELSR2^+^ cells were found to have increased expression of *ZP3*, *HLA-DOB*, *HLA-DQA2*, *HLA-DQA1* and *IL9R*, which were enriched in the Immunoglobulin Mediated Immune Response biological process, and *IL1B*, *S100A9*, *S100A8* enriched in the Leukocyte Aggregation pathway (**Supplemental Data 6**). In contrast, none of the genes upregulated in Cluster 1 – Differentiating T_EM_ in HC_NR participants were enriched in any biological processes. In line with our findings of an elevated cytotoxic signature in CELSR2+ HC_NR cells, we found that the HC_NR CELSR2+ cells from both Cluster 4 – Cytotoxic T_EM_ and Cluster 6 – Activated T_EM_ expressed higher levels of genes enriched in the Natural Killer Cell Mediated Immunity biological process (**Supplemental Data 7**).

Overall, despite a higher frequency of CESLR2^+^ in CD4 T_EM_ in PD_R, transcriptomic changes compared to CELSR2^-^ are minor.

### PD_R have decreased frequency of Granulysin^+^ cytotoxic CD4 T_EM_ cells

The most striking difference we discovered is the almost complete lack of Cluster 4 – Cytotoxic T_EM_ in PD_R compared to HC_NR. As cytotoxic CD4 T cells have been previously described in PBMC from PD participants^22, 23^, we next set out to validate our scRNAseq finding using flow cytometry. To address this, we first extracted the gene signature of cluster 4 to identify cytotoxic T cells by flow cytometry and retained *granzyme B*, *perforin* and *granulysin* as specific markers (**Figure 4A, Supplemental Figure 6A**). We then quantified the total population of cytotoxic CD4 T cells, either identified by the double expression of Granzyme B and Perforin, or single expression of Granulysin in HC_NR (n=11; Female=9, Male=2) and PD_R (n=8; Female=0, Male=8, p=0.001) study participants (**Supplemental Data 8**). Both populations were distributed in a similar fashion in HC_NR and PD_R, both in regards to frequency of expressing cells (GranzymeB^+^Perforin^+^, p-value=0.091 and Granulysin^+^, p-value=0.492) (**Figure 4B, 4C**), and MFI of Granulysin (p=0.129), perforin (p=0.152), and granzyme B (p=0.272; **Supplemental Figure 6B**). After we confirmed that there were no bias in the distribution of memory and naive subpopulations (**Supplemental Figure 6C**), we determined the frequency of T_EM_ within GranzymeB^+^Perforin^+^ (**Figure 4D**) or Granulysin^+^ (**Figure 4E**). As in the scRNAseq dataset, we found a significantly lower frequency of Granulysin^+^ cytotoxic CD4 T_EM_ cells in PD_R compared to HC_NR (p-value=0.009). Importantly, this depletion was specific for CD4 T cells, as no difference was seen in frequency of GranzymeB^+^Perforin^+^ or Granulysin^+^ cells among all CD8 T cells, or of T_EM_ within the GranzymeB^+^Perforin^+^ or Granulysin^+^cytotoxic CD8 population (**Supplemental Figure 6D, 6E**).

**Figure 4.**
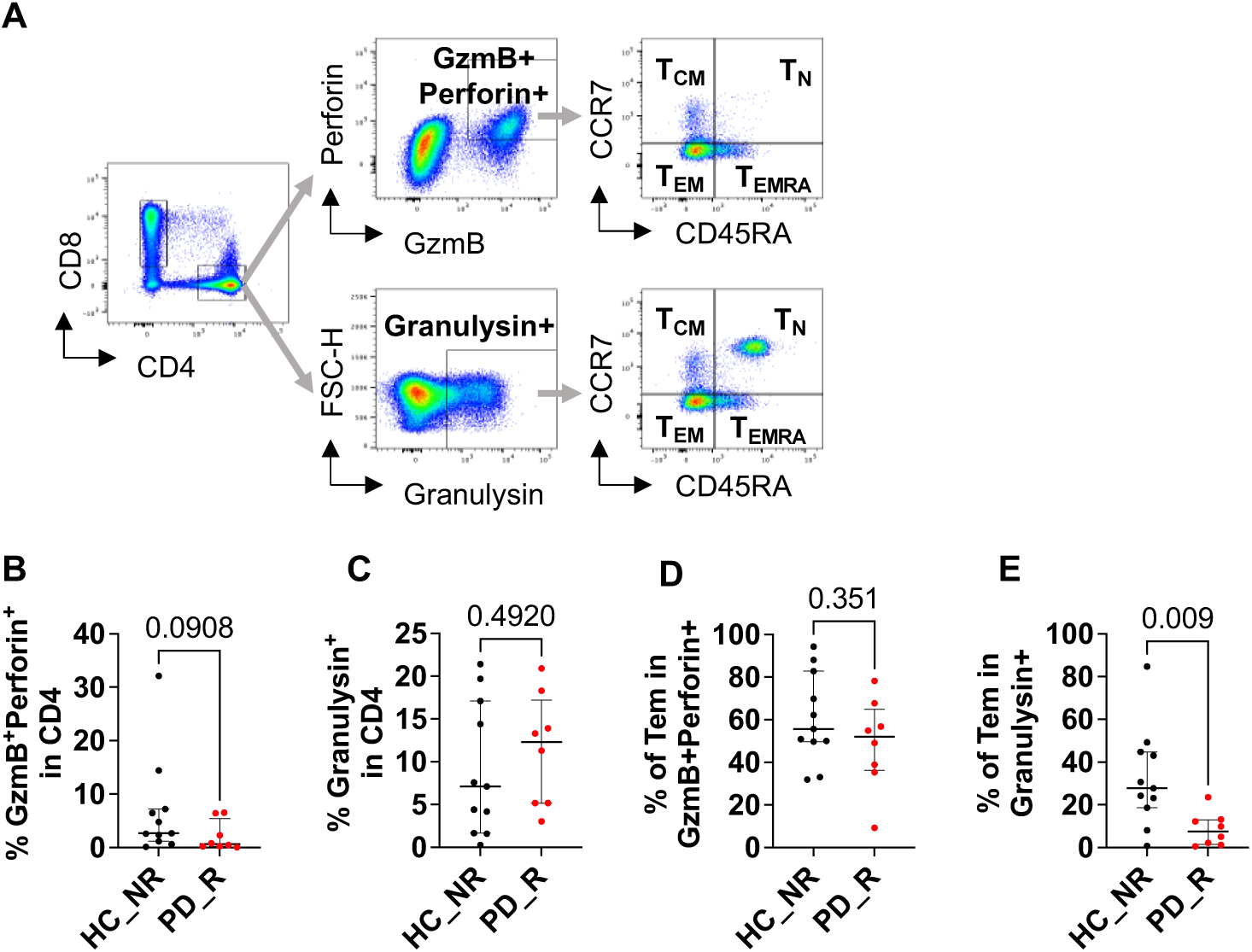
Cytotoxic T cells from PD_R are differentially enriched in T_EM_ than HC_NR. **(A)** Representative gating strategy for the identification of CD4 T cells expressing both Granzyme B (GzmB) and perforin, or Granulysin, and the further memory cell phenotyping of the cytotoxic cell subsets, in HC_NR (n=11) and PD_R (n=8) participants. Complete gating strategy is displayed in **Supplemental** Figure 5. The frequencies of cytotoxic CD4 T cells identified by the double expression of GzmB and perforin **(B)** or Granulysin **(C)** were analyzed. Frequencies of T_EM_ (CD45RA^-^CCR7^-^) within GzmB^+^Perforin^+^ **(D)** and Granulysin^+^ **(E)** were analyzed. Results are represented as median with interquartile range. Mann Whitney U test.

## Discussion

In this study, we report for the first time that CESLR2 protein is expressed by T cells. We show a higher frequency of CELSR2 positive cells in CD4 T_EM_ from PD participants responding to *α*-syn. Using scRNA-seq on sorted CD4 T_EM_ CELSR2^+^ and CELSR2^-^ populations from PD_R and HC_NR, we discovered that PD_R CELSR2^+^ cells are enriched in cells with transcriptional profiles involved in T cell activation and differentiation, and is depleted in cytotoxic cells. Using flow cytometry, we validated that PD_R exhibit a depletion of Granulysin^+^ cytotoxic CD4 T_EM_ cells.

ScRNAseq analysis revealed cluster distributions differences dependent on both CELSR2 expression and disease status. While Cluster 6 – Activated T_EM_ were increased in both CELSR2^+^ samples, the increased and decreased frequency of Cluster 1 – Differentiating T_EM_ and Cluster 4 – Cytotoxic T_EM_, respectively, appeared to be related to disease status rather than CELSR2 expression. Cluster 1 – Differentiating T_EM_ was enriched in PD_R compared to HC_NR, with the highest frequency within the CELSR2^+^ samples. The high frequency of these cells makes them of interest for future studies of immunopathology in PD pathology. These cells express high levels of genes involved in interactions with antigen presenting cells. It will therefore be of interest to study if the increased frequency of these cells in PD participants contributes to the previously reported hyperactivation of glial cells^24^, the CNS-resident antigen presenting cells. Our most striking discovery is the almost complete absence of Cluster 4 – Cytotoxic T_EM_ in PD_R compared to HC_NR. Using flow cytometry, we verified that Granulysin^+^ cytotoxic CD4 T cells are depleted among PD_R T_EM_. As T_EM_ rapidly migrate into inflamed tissues upon activation^25^, it will be of great interest to further study if the depletion of the Granulysin^+^ cytotoxic CD4 T_EM_ cells has previously occurred due to an increased egress of these cells into peripheral tissues such as the CNS. Previous studies have reported increased frequencies of CD4 T cells in the brain of PD participants^26^, but the phenotype and specificity of these cells still remain largely unknown. As we in this study focused on PD_R, and therefore did not include PD_NR participants, future studies should further validate if the observed alterations in proportion of T cell subpopulations is specific to PD_R or represent a general alteration of the T cell compartment in people living with PD. However, as *α*-syn responses are strongest close to diagnosis^14^, these findings from PD_R are most likely representative of early PD pathology.

CESLR2 is a member of the set of atypical cadherins crucial during embryonic development due to its involvement in the planal cell polarity^27–29^. In human, biallelic *CELSR2* mutations lead to the development of Joubert Syndrome, which is characterized by diverse hindbrain and midbrain anomalies^30^, which often clinically manifests as hypotonia, development delay and subsequent ataxia and apraxia of speech. Other roles for CELSR2 include axon pathfinding and brain wiring^31^, but to our knowledge has never been associated to T cell biology. When we investigated the selected expression of CELSR2 on T cells, we highlighted its higher expression within the memory, and T_EM_ subpopulations specifically. This can suggest its increased expression is linked to antigen stimulation, emphasized on PD participants that respond to *α*-syn. Previous studies have shown that CELSR2 promote activation of macrophages and Schwann cells through the activation of the Wnt/β-catenin and RAS-ERK pathways^32, 33^, and to inhibit cell death by increasing expression of antioxidant enzymes^34^. In line with these reports, scRNAseq analysis of sorted CD4 T_EM_ CELSR2^+^ and CELSR2^-^ revealed a higher proportion of Cluster 1 – Differentiating T_EM_ and Cluster 6 – Activated T_EM_ in CELSR2^+^ population in both PD_R and HC_NR. It is therefore possible that the increased frequency of CELSR2 expressing cells in PD_R compared to HC_NR reflects an increased activation of T cells in PD^35^. This is in line with previous reports of inflammation playing a fundamental role in PD pathology^35, 36^. Furthermore, in Cluster 4 – Cytotoxic T_EM_ within HC_NR, the cytotoxic gene signature is enriched within the CELSR2^+^ subpopulations. As cytotoxic T cells are associated with chronic stimulation and a memory phenotype^37^, it supports our hypothesis that *CELSR2* expression is induced by TCR stimulation. Our study focused on CESLR2 as its expression is reliable and higher on T cells from PD_R than HC_NR. The identification of CELSR2 was based on a protein coding gene list extracted from a bulk RNAseq on sorted CD4/CD8 T mem^15^, which was narrowed down to the availability of a commercial antibody. Thus, other potentially relevant targets can be found in the gene list. While the transcriptome of CELSR2 expression was not associated with major changes in CD4 T_EM_, other membrane proteins could be specific to PD_R T cells and thus be used to develop targeted therapies for PD patients reactive to *α*-syn. As we discovered a differential CELSR2 expression on CD4 T_EM_ in PD_R compared to HC_NR, it led us to investigate the heterogeneity using scRNAseq. However, focusing on T_EM_ limited us from discovering other potential differences between HC_NR and PD_R within the total CD4 T cell population. As our study is limited by the low number of samples available from female PD study participants, and biological sex differences has been reported in the proportions of T cell subpopulations^38^, the findings in this study should be further validated in a larger sex-balanced cohort of donors.

To conclude, we showed for the first time that CELSR2 is expressed by T cells, enriched in CD4 T_EM_, and greater in PD_R. Using scRNAseq and flow cytometry, we discovered a differential memory depletion of cytotoxic CD4 T cells in PD_R, which represents an opportunity for a better comprehension of PD pathology and subsequent therapeutic opportunities.

## Methods

### Study approval

All participants provided written informed consent for participation in the study. Ethical approval was obtained from the Institutional Review Boards at La Jolla Institute for Immunology (LJI; Protocol Nos: VD-124 and VD-118), Columbia University Irving Medical Center (CUIMC; protocol number IRB-AAAQ9714 and AAAS1669), Rush University Medical Center (RUMC; Office of Research Affairs No.16042107-IRB01), University of California San Diego (UCSD; protocol number 161224), the University of Alabama at Birmingham (UAB; protocol number IRB-300001297) and Shirley Ryan AbilityLab/Northwestern University (protocol number STU00209668-MOD0005).

### Study participants

Subjects with idiopathic PD and HCs were recruited by the Movement Disorders Clinic at the Department of Neurology at CUIMC, by the clinical core at LJI, by the Parkinson and Other Movement Disorder Center at UCSD, by Parkinson’s disease and Movement Disorders Care at RUMC, by movement disorder specialists at UAB Movement Disorders Clinic and by the movement disorder specialists at the Parkinson’s disease and Movement Disorders program at Shirley Ryan AbilityLab. Inclusion criteria for PD patients consisted of i) clinically diagnosed PD with the presence of bradykinesia and either resting tremor or rigidity ii) PD diagnosis between ages 35-80 iii) history establishing dopaminergic medication benefit, iv) ability to provide informed consent. Exclusion criteria for PD were atypical parkinsonism or other neurological disorders, history of cancer within past 3 years, autoimmune disease, and chronic immune modulatory therapy. Age-matched HC were selected on the basis of i) age 45-85 and ii) ability to provide informed consent. Exclusion criteria for HC were the same as PD except for the addition of self-reported PD genetic risk factors (i.e., PD in first-degree blood relative). For the LJI cohort, PD was self-reported. Individuals with PD recruited at CUIMC, UCSD, and Shirley Ryan AbilityLab all met the UK Parkinson’s Disease Society Brain Bank criteria for PD. Cohort demographics are listed in **Supplemental Data 1** (protein target screening participants), and **Supplemental Data 2** (scRNAseq participants**), and Supplemental Data 8** (cytotoxic T cell analysis participants).

### PBMC isolation

Venous blood was collected from each participant in either heparin or EDTA containing blood bags or tubes. Blood samples not collected at LJI were shipped overnight at 4°C to LJI. PBMC were isolated from whole blood by density gradient centrifugation using Ficoll-Paque plus (GE #17144003). In brief, blood was first spun at 1850 rpm for 15 min without brake to remove plasma. Plasma depleted blood was then diluted with RPMI, and 35 mL of blood was carefully layered on tubes containing 15 mL Ficoll-Paque plus. These tubes were then centrifuged at 1850 rpm for 25 min with the brakes off. The interphase cell layer resulting from this spin were collected, washed with RPMI twice at 1850 rpm for 10 min with a low brake, platelets were removed by centrifugating at 800 rpm for 10 min without brake. Cells were then counted, and cryopreserved in 90% v/v FBS and 10% v/v dimethyl sulfoxide and stored in liquid nitrogen until tested. The detailed protocol for PBMC isolation can be found at protocols.io^39^. Cryopreserved PBMC were thawed by placing the vial in 37°C water bath for 1 min, and washing in medium supplemented RPMI (Corning) supplemented with 5 % human serum (Gemini Bio-Products), 1 % Glutamax (Gibco), 1 % penicillin/streptomycin (Omega Scientific) in the presence of benzonase (20 μL/10 mL). The detailed protocol for PBMC isolation can be found at protocols.io (https://dx.doi.org/10.17504/protocols.io.bw2ipgce).

### Flow cytometry

*Antibody screening*. PBMC were plated in a U-bottom 96 plate at a density of 10^6^ cells/well. Cells were stained using a Fixable Viability Dye eF506 (Thermo Fisher) and Fc receptors were blocked (BD Biosciences) before membrane staining with antibodies for 20 min at 4°C. Membrane staining was performed sequentially as tested antibodies are purified; first purified antibodies, then a PE-coupled secondary antibody, and finally the remaining coupled antibodies. Stained cells were washed twice and acquired on a LSR II or Fortessa X-20 (BD Biosciences). FlowJo software (v10.10.0, Tree Star) was used for the analysis. Gating strategy is shown in **Supplemental Figure 1A**. The detailed protocol can be found at DOI: dx.doi.org/10.17504/protocols.io.rm7vzk934vx1/v1 (Private link for reviewers: https://www.protocols.io/private/A0A48C69E7DA11EFBF9E0A58A9FEAC02 to be removed before publication).

*scRNAseq.* 20×10^6^ cells per participant were used. Fc receptors were blocked (BD Biosciences) before membrane staining with antibodies and a specific hashtag (TotalSeq B; Biolegend) for 20 min at 4°C. Cells were washed and DAPI was added to stain dead cells. Two populations were sorted using a BD FACSAria II (BD Biosciences): DAPI^-^CD3^+^CD4^+^CD8^-^CD45RA^-^CCR7^-^ CESLR2^+^ and DAPI^-^CD3^+^CD4^+^CD8^-^CD45RA^-^CCR7^-^CESLR2^-^. Gating strategy is shown in **Supplemental Figure 1E**. The detailed protocol can be found at DOI: dx.doi.org/10.17504/protocols.io.q26g7mn48gwz/v1 (Private link for reviewers: https://www.protocols.io/private/37159022E7DF11EFB4270A58A9FEAC02 to be removed before publication.)

*Cytotoxic flow panel.* 4×10^6^ cells per participant were stimulated with 1ug/mL PMA, 1μg/mL ionomycin in the presence of GolgiPlus and GolgiStop (BD Biosciences) for 4h at 37°C 5% CO_2_. 4×10^6^ cells per participant remained unstimulated and were only incubated with GolgiPlus and GolgiStop (BD Biosciences) for 4h at 37°C 5% CO_2_. Cells were washed and cells were stained using a Fixable Viability Dye eF506 (Thermo Fisher) and Fc receptors were blocked (BD Biosciences) for 20 min at 4°C, before membrane staining with antibodies for 30 min at 4°C. For intracellular staining, cells were permeabilized 30 min at RT with Intracellular Fixation & Permeabilization Buffer Set (Thermo Fisher), stained for 45 min with antibodies at room temperature and then fixed with 4% PFA. Stained cells were washed twice and acquired on a Fortessa X-20 (BD Biosciences). FlowJo software was used for the analysis. Gating strategy is shown in **Supplemental Figure 6A**. All antibodies used are listed in **Supplemental Data 9**. The detailed protocol can be found at DOI: dx.doi.org/10.17504/protocols.io.yxmvm9bn9l3p/v1 (Private link for reviewers: https://www.protocols.io/private/2904289CE7E511EFBF9E0A58A9FEAC02 to be removed before publication).

### scRNA-Seq Library Preparation and Sequencing

A total of 16 unique samples were hashed using BioLegend Total-Seq B antibodies. Cells were sorted and approximately 12,000 cells/sample were targeted and loaded on the 10X Chromium Controller (10X Genomics). We used the Chromium Next GEM Single Cell 3’ HT Reagent Kits v3.1 (Dual Index) with Feature Barcode technology for Cell Surface Protein and Cell Multiplexing and performed cDNA synthesis, amplification, and library preparation for gene expression and hashtag antibodies according to manufacturer’s protocol. cDNA and final libraries were quality checked for fragment size by capillary electrophoresis using the Tapestation 4200 (Agilent), and quantified using the Qubit Flex assay (Thermo Fisher), before pooling and sequencing on a NovaSeq 6000 S4 v1.5 Flowcell (Illumina) to a depth of > 20,000 reads/cell for gene expression and > 5,000 reads/cell for hashtag antibody libraries. The detailed protocol can be found at DOI: dx.doi.org/10.17504/protocols.io.8epv5n32nv1b/v1 (Private link for reviewers: https://www.protocols.io/private/C5F448C61BCC11F0ABFF0A58A9FEAC02 to be removed before publication).

### Single-cell RNAseq analysis

The CellRanger (v7.1) software suite was used to convert Illumina sequencer’s base call files to FASTQ files, to align sequencing reads to the GRCh38 human reference genome and generate count matrix, and to demultiplex samples based on the TotalSeq B hashtag sequences. Downstream analyses were performed in R (v4.3.2) using the Seurat package (v5.1)^40–44^. Cells identified as doublets during hashtag demultiplexing, or had too high mitochondrial gene expression (>6%), too low number of detected genes (<1000), too high number of detected genes (>3500), too low Unique molecular identifier (UMI) count (<2000), or too high UMI count (>10000) were excluded as part of the standard quality control process. The SCTransform package (v0.4.1), using the glmGamPoi method (v1.14.3), was used to normalize the samples and regress out the percentage of mitochondrial genes. The samples where thereafter integrated using the IntegrateData function, using predefined features and anchors defined using the SelectIntegrationFeatures and FindIntegrationAnchors functions, respectively. The SCT normalization methods was used, and dimensions 1 to 30 for the anchor weighting procedure. A principal component analysis was thereafter performed using the RunPCA function and the top 30 dimensions were used for the RunUMAP function. To identify clusters, we next used the FindNeighbors function using the top 30 PCA dimensions and a k.param of 200, and finally used the FindClusters function with a resolution of 0.5. The optimal resolution was identified using the Clustree package (0.5.1). Cluster markers were defined using the FindAllMarkers function using the SCT assay, minimum percentage of cells in the cluster expressing a gene being 25% and a log fold change cut off at 0.25.

### Differential gene expression and biological process enrichment analyses

Differentially expressed genes were identified using the FindMarkers function using the MAST algorithm (v1.28.0) on the SCT integrated expression data^45^, including only genes expressed in at least 1% of the cells in either group. Genes were defined as differentially expressed if it had an adjusted p value <0.05 and a log_2_ fold change > 1. Enrichment of differentially expressed genes in biological processes were identified using the DEenrichRPlot function and the GO_Biological_Process_2023 enrichR library of genes. All plots were produced using Seurat or ggplot2 (3.5.1). Module scores were created using the AddModuleScore function. The Leukocyte mediated cytotoxicity module score was based on 109 genes both detected in our samples as well included in the Leukocyte mediated cytotoxicity biological process (GO:0001909), and the Ribosomal gene signature was based on the ribosomal genes identified as markers of Cluster 0 – RP^+^ T_EM_ (**Supplemental Data 3**).

## Statistical Analysis

Data analysis and statistical comparisons were conducted using GraphPad Prism version 10.2.3 and R. We used a two-tailed Mann-Whitney U test or 2way ANOVA and Tukey’s multiple comparisons test when analyzing CELSR2, cytokines, memory subpopulations and surface marker expressions comparing 2 groups with 1 or 2 variables. A one sample t and Wilcoxon test was used for the fold increase analysis. The Fisher’s exact test was used to compare proportion of male and female study participants. Paired or unpaired t tests were used to compare frequency of scRNAseq clusters within PD_R participants or between PD_R and HC_NR participants, respectively. All statistical tests, sample sizes and error bar descriptions of graphs are detailed in the legends of respective figures.

## Data availability

Raw and processed scRNAseq data is available through the National Center of Biotechnology Information’s Gene Expression Omnibus (GEO) at the accession number GSE289241. Source data for all figures are provided with this paper, and links to raw data is provided in the Key resource table.

## Code availability

Code used to analyze scRNAseq data has been deposited to Zenodo, as described in the Key resource table.

## Supporting information

Key Resource Table

## Acknowledgments

We would like to thank all the participants for donating samples for this study. We are also grateful to the clinical core and lab members for processing the blood samples, the sequencing core facility for help with single-cell RNAseq library prep and sequencing (RRID:SCR_023107), and the flow cytometry core for cell sorting at La Jolla Institute for Immunology (RRID:SCR_014832). The NovaSeq 6000 was acquired through the Shared Instrumentation Grant (SIG) Program (S10) S10OD025052.

This work was supported by Aligning Science Across Parkinson’s (ASAP-000375 to C.L.A and D.S.) through the Michael J. Fox Foundation for Parkinson’s Research (MJFF), by Kyowa Kirin, Inc. (KKNA-Kyowa Kirin North America), and by the Swedish Research Council (to E.J. grant references 2024-00175). For open access, the authors have applied a CC-BY public copyright license to all author-accepted manuscripts arising from this submission. The funders had no role in study design, data collection, analysis, publication decision, or manuscript preparation.

## Author Contributions

A.F. performed the experiments. A.F. and E.J. analyzed the data. A.Fra, I.L., J.G.G., and R.N.A. recruited participants and performed clinical evaluations. A.F, E.J, C.S.L.A., and A.S. wrote the paper with substantial edits by D.S. and other authors. A.F., E.J., C.S.L.A., and A.S. designed and discussed the data. All authors read, edited, and approved the paper.

