## Supplemental Information for "Differential memory enrichment of cytotoxic CD4 T cells in Parkinson’s disease patients reactive to α-synuclein"

**Supplementary information**

**Data**

**Supplementary Data 1. Demographics of donors used for the target screening.** P-value reported for comparison of gender proportion was obtained using a Fisher's exact test, and two-tailed Mann-Whitney U test for the comparison of age.

**Supplementary Data 2. Demographics of donors used for the single cell RNA sequencing experiment.** P-value reported for comparison of gender proportion was obtained using a Fisher's exact test, and two-tailed Mann-Whitney U test for the comparison of age.

**Supplementary Data 3. Top 50 genes for each cluster**

**Supplementary Data 4. Cluster 1 PD_R CELSR2+ vs PD_R CELSR2- DEG enrichment analysis**

**Supplementary Data 5. Leukocyte mediated cytotoxicity module score genes**

**Supplementary Data 6. Cluster 1 PD_R CELSR2+ vs HC_NR CELSR2+ DEG enrichment analysis**

**Supplementary Data 7. Cluster 4 and 6 PD_R CELSR2+ vs HC_NR CELSR2+ DEG enrichment analysis**

**Supplementary Data 8. Demographics of donors used for the CD4 cytotoxic analysis.** P-value reported for comparison of gender proportion was obtained using a Fisher's exact test, and two-tailed Mann-Whitney U test for the comparison of age.

**Supplementary Data 9. Antibodies used for the study**

### Figures

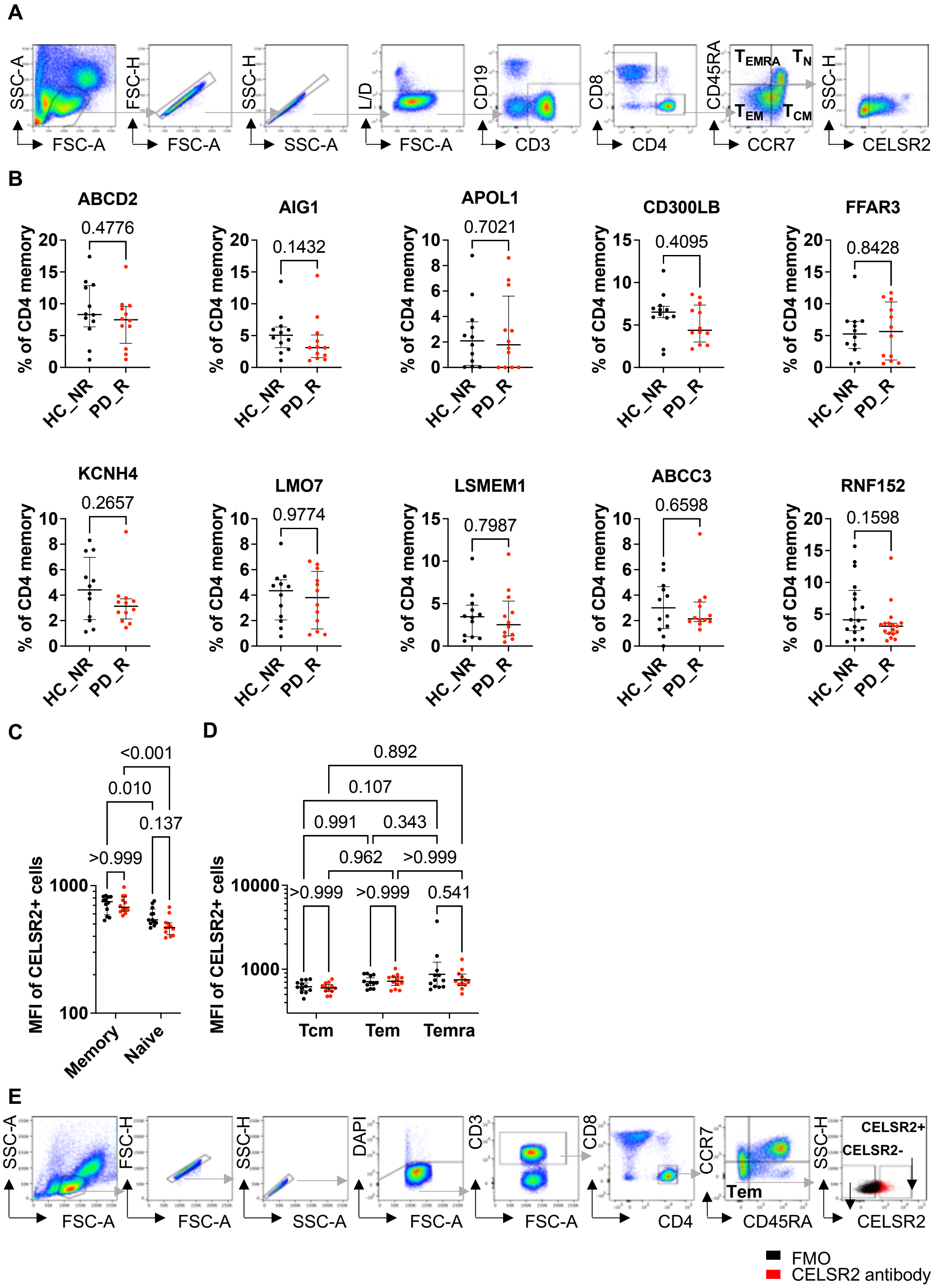


**Supplemental Figure 1. Gating strategies.** (**A**) Representative gating strategy to identify CD4 T cell naïve ($T_N$, $CD45RA^+CCR7^+$), central memory ($T_{CM}$, $CD45RA^-CCR7^+$), effector memory

($T_{EM}$, CD45RA$^{-}$CCR7$^{-}$) and effector memory cells re-expressing CD45RA ($T_{EMRA}$, CD45RA$^{+}$CCR7$^{-}$) **(B)** Expression of all markers was analyzed on total memory CD4 T cells (CD45RA$^{+}$CCR7$^{-}$, CD45RA$^{-}$CCR7$^{+}$, and CD45RA$^{-}$CCR7$^{-}$ cells) in PD_R (n=12) and HC_NR (n=12). Geometric mean fluorescent intensity (MFI) of CELSR2 staining on $T_N$ and total memory ($T_{CM}$, $T_{EM}$, and $T_{EMRA}$) (**C**), or within $T_{CM}$, $T_{EM}$, and $T_{EMRA}$ cells (**D**). **(E)** Cell sorting of CD4 $T_{EM}$ CELSR2$^{+}$ and CELSR2$^{-}$ was performed on PBMCs from 4 HC_NR and 4 PD_R. Cells were first gated on lymphocytes, doublets exclusion, then in DAPI$^{-}$CD3$^{+}$CD4$^{+}$CD8$^{-}$CD45RA$^{-}$CCR7$^{-}$, both CESLR2$^{+}$ and CELSR2$^{-}$ subpopulations were sorted. FMO were performed as negative population controls (black). Results are represented as median with interquartile range. Mann Whitney U test was used to compare between two groups, and a 2way ANOVA and Sidak's multiple comparisons test when comparing multiple groups.

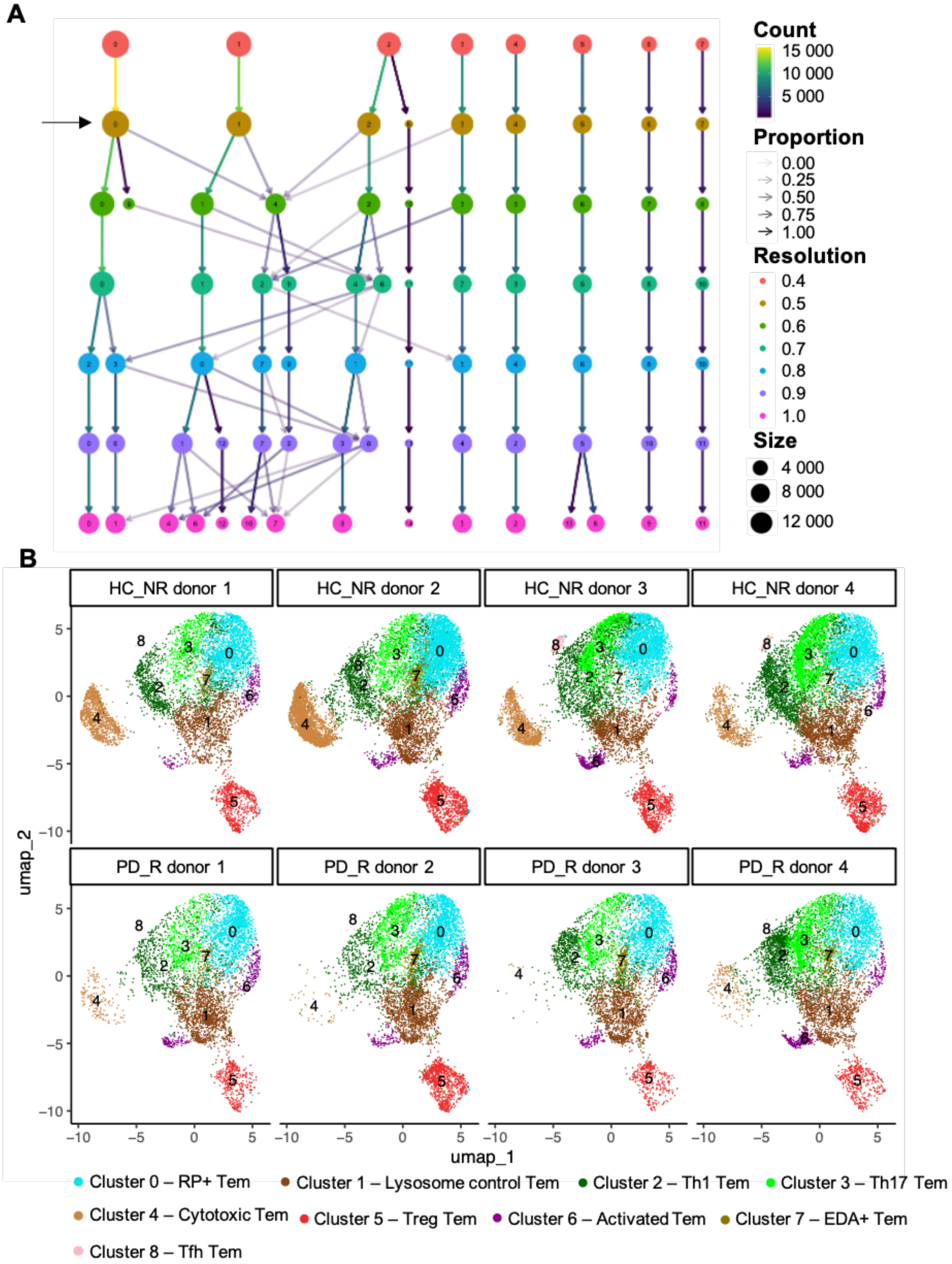


**Supplemental Figure 2. Clustering tree and scRNAseq UMAPs per participant. (A)** The impact of cluster resolution on the number of identified clusters visualized using the Clustree package. **(B)** Individual UMAP plots for each HC_NR and PD_R participant.

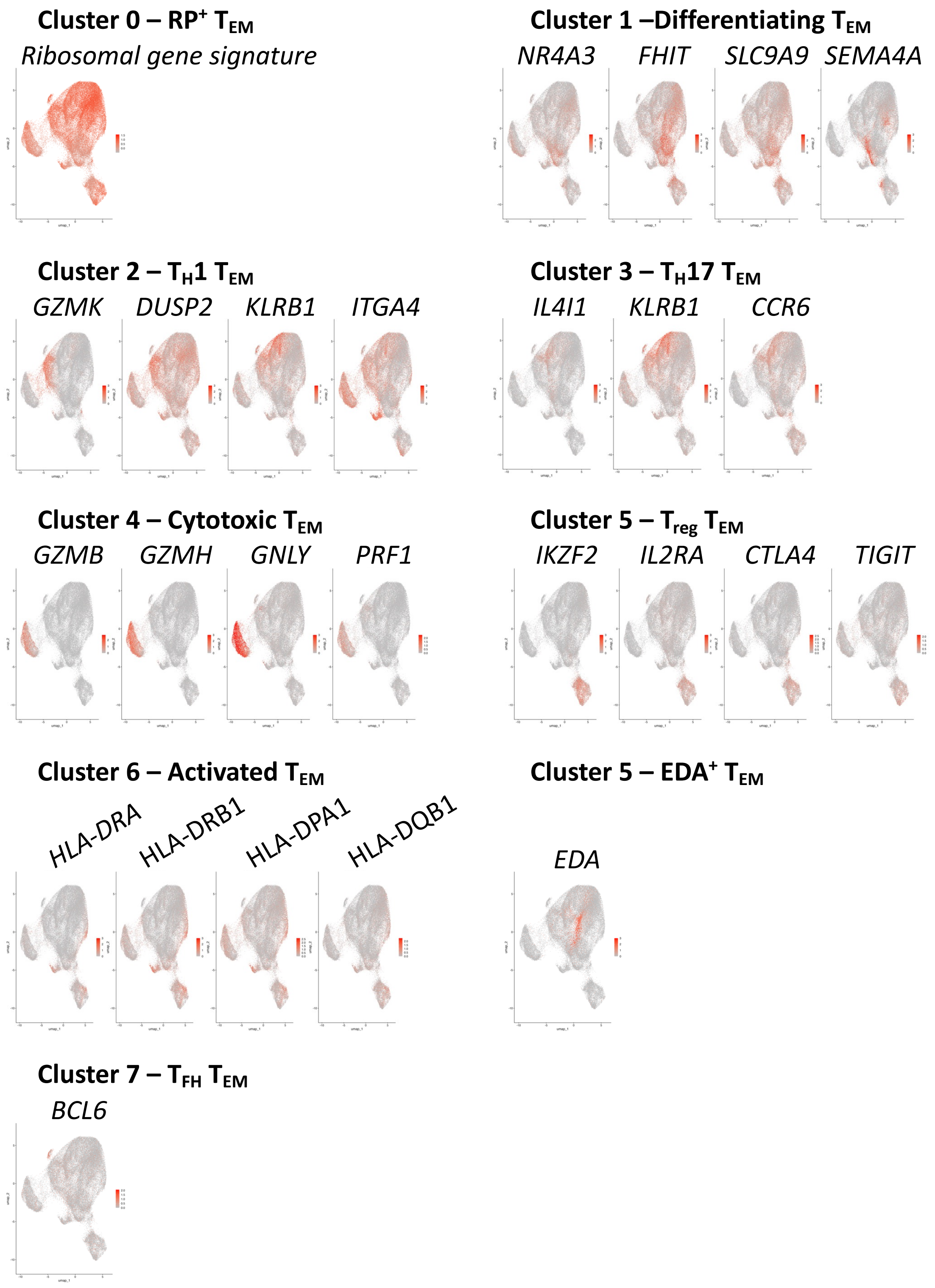


**Supplemental Figure 3. UMAP plots of cluster markers.** Expression of markers used to classify cluster are projected on UMAP plots. Remaining makers can be found in **Supplemental Data 3**.

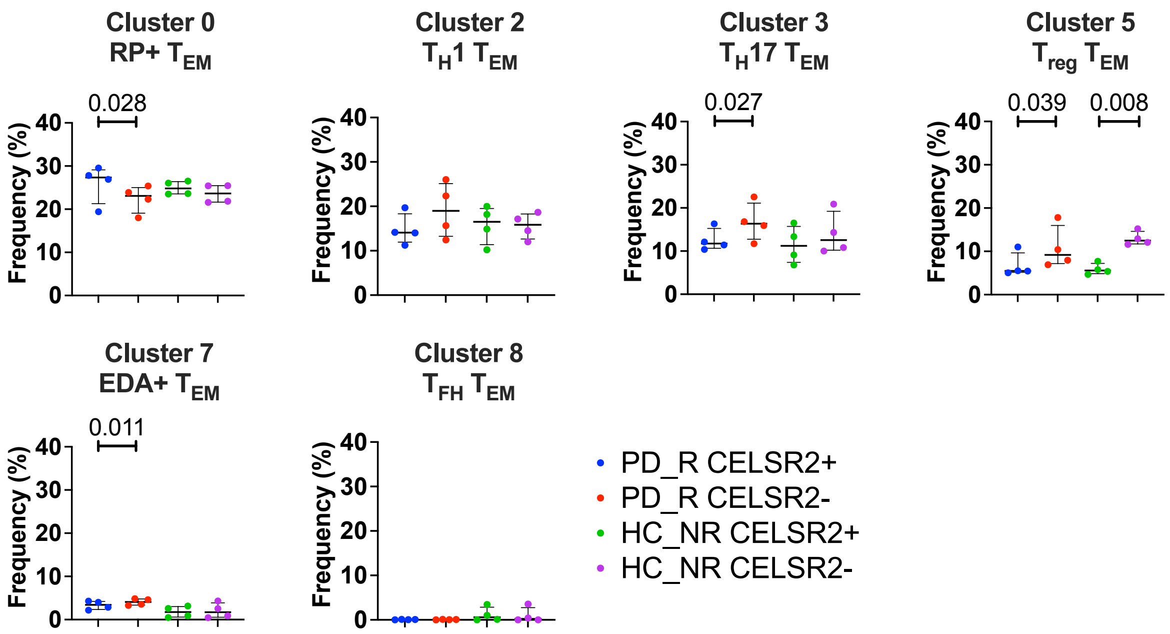


**Supplemental Figure 4. Cluster frequency for cluster found not to differ between PD_R and HC_NR samples.** The frequency of Clusters 0, 2, 3, 5, 7 and 8 represented as median with interquartile range. Frequencies were compared using paired t test within PD_R or HC_NR samples, and unpaired t test to compare frequency between PD_R and HC_NR subgroups.

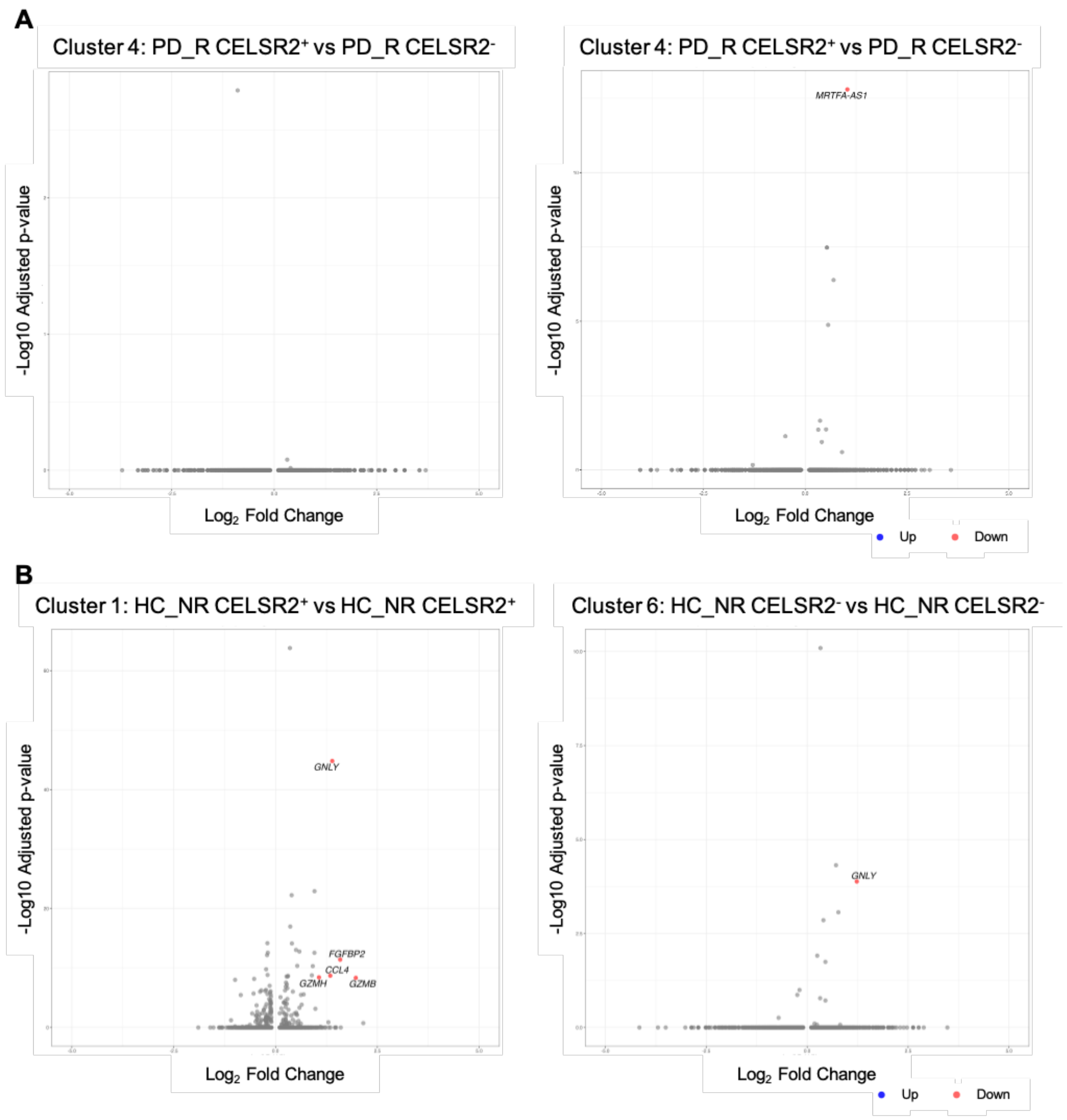


**Supplemental Figure 5. Expression of CELSR2 does not significantly impact transcriptomics in PD_R participants in Cluster 4 – Cytotoxic $T_{EM}$ and Cluster 6 – Activated $T_{EM}$, or for HC_NR participants in Cluster 1 – Differentiating $T_{EM}$ and Cluster 6 – Activated $T_{EM}$.** Volcano plots comparing differential gene expression of PD_R CELSR2$^{+}$ vs PD_R CELSR2$^{-}$ in Cluster 4 – Cytotoxic $T_{EM}$ **(A)** and Cluster 6 – Activated $T_{EM}$ (**B**), and HC_NR CELSR2$^{+}$ vs HC_NR CELSR2$^{-}$ in Cluster 1 – Differentiating $T_{EM}$ (**C**) and Cluster 6 – Activated $T_{EM}$ (**D**). Dots colored in red are significantly upregulated in CELSR2+ cells (adjusted p value < 0.05 and $Log_2$ fold change >1).

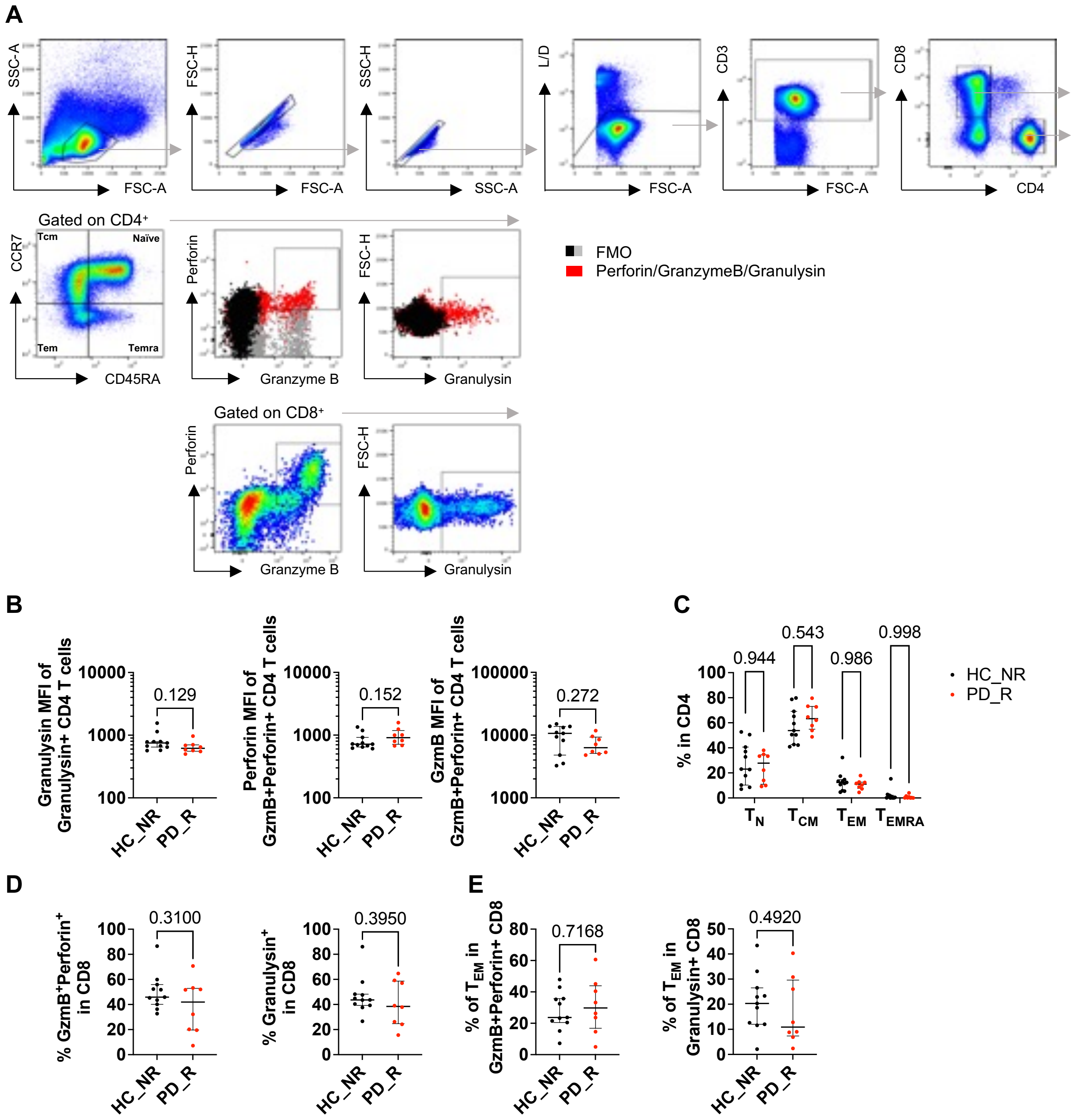


**Supplemental Figure 6. Cytotoxic T cell gating strategy and analysis. (A)** Gating strategy depicting the analysis of cytotoxic CD4 T cells in HC_NR (n=11) and PD_R (n=8). PBMCs were first gated on lymphocytes, then doublets exclusion and L/D$^-$CD3$^+$. Further gating on CD4$^+$CD8$^-$ to analyze memory T cell subpopulations ($T_{CM}$, $T_{EM}$ and $T_{EMRA}$) and naïve as well as GranzymeB$^+$(GzmB)Perforin$^+$ double expressing and Granulysin$^+$ cells. FMO were performed as negative controls (black/grey). Expressions of Granzyme B, Perforin and Granulysin in $CD4^{-}CD8^{+}$ were used as positive controls. **(B)** The geometric mean fluorescent intensity (MFI) of Granulysin on Granulysin+ CD4 T cells (left panel), as well as Perforin (middle panel) and GzmB (right panel) MFI on $GzmB^{+}Perforin^{+}$ CD4 T cells was quantified. (**C**) Memory subpopulations ($T_{CM}$, $T_{EM}$ and $T_{EMRA}$) and naïve cells distributions within CD4 T cells. **(D)**. Frequency of CD8 T cells co-expressing GzmB and perforin (left panel), or Granulysin (right panel). (**E**) Frequency of $T_{EM}$ among GzmB+Perforin+ or Granulysin+ CD8 T cells. P-value was obtained using a 2way ANOVA and Sidak's multiple comparisons test when comparing multiple groups, and a non-paired Mann Whitney U test. Results are represented as median with interquartile range.
